# AlfaDAX-Derived ActRIIA/B Antibody with Semaglutide Enhances Fat Loss and Improves Weight-Loss Quality in DIO Mice

**DOI:** 10.64898/2026.07.09.737400

**Authors:** Ning Zhang, Yongkang Long, Zhangzhan Xu, Gaojian Chen, Ao Wang, Weihai Chen, Zhongwei Chen, Zhiming Liang, Kingsley Leung, Liang Chen

## Abstract

GLP-1 receptor agonists achieve weight loss but are associated with clinically significant reductions in lean mass. Activin type II receptors (ActRIIA and ActRIIB) mediate signaling of myostatin and activin A, both of which negatively regulate muscle growth, suggesting that dual blockade of these receptors may preserve or increase lean mass while promoting fat loss. In this study, we developed anti-ActRIIA/B antibodies using AI-driven platforms (AlfaDAX) and selected the lead candidate AB130-165 based on *in vitro* binding, functional blocking, and developability assessments. Compared with a laboratory-prepared bimagrumab analog, AB130-165 exhibited potent dual inhibition of ActRIIA/B signaling, with a 9.5-fold higher functional blocking activity against activin A-induced SMAD signaling and 1054-fold improvements in binding affinity for ActRIIA (K_D_ = 0.204 pM), 10-fold for ActRIIB (K_D_ = 0.243 pM), respectively. In diet-induced obese mice, combination therapy with AB130-165 and semaglutide resulted in a 33.4% body weight reduction, which was superior to semaglutide monotherapy (−24.3%) and the bimagrumab combination group (−25.5%). Moreover, the combination significantly improved body composition, reducing fat mass percentage by 77.8% (*vs*. 65.0% in the bimagrumab combination group) and increasing the lean-to-body weight ratio to 67.3% (*vs*. 62.3%), demonstrating superior fat loss with better preservation of lean mass. Collectively, these findings establish AB130-165 as a differentiated anti-ActRII antibody that enables high-quality weight loss, and its combination with semaglutide shows superior efficacy over bimagrumab-based regimens. With favorable developability and potential for long-acting subcutaneous administration, AB130-165 represents a promising next-generation therapeutic candidate for obesity and muscle-sparing weight management.

## 1 Introduction

The global obesity pandemic has reached unprecedented proportions, with over 650 million adults currently classified as obese, significantly increasing the risk of type 2 diabetes (T2DM), cardiovascular disease, and metabolic dysfunction-associated steatohepatitis (MASH)^[1, 2]^. The recent clinical success of Glucagon-Like Peptide-1 receptor agonists (GLP-1RAs), such as semaglutide and tirzepatide, has transformed the therapeutic landscape, demonstrating weight loss efficacy previously achievable only through bariatric surgery^[3, 4]^. However, a significant clinical concern has emerged: a substantial portion (up to 40%) of the weight lost through GLP-1RA therapy is derived from lean body mass rather than adipose tissue^[5–8]^.

The loss of lean mass, particularly skeletal muscle, is metabolically deleterious. Skeletal muscle is the primary site for glucose disposal and a major determinant of resting metabolic rate (RMR)^[9, 10]^. Excessive muscle loss can lead to “sarcopenic obesity,” potentially predisposing patients to functional decline, reduced insulin sensitivity, and a heightened risk of weight regain upon treatment cessation^[11]^. Consequently, the next frontier in obesity management is shifting from “quantity of weight loss” to “quality of weight loss”—defined as the selective reduction of fat mass while preserving or enhancing lean mass.

The Transforming Growth Factor-beta (TGF-β) superfamily, specifically the signaling pathways mediated by Activin type II receptors (ActRIIA and ActRIIB), plays a fundamental role in regulating muscle homeostasis^[12, 13]^. Two key ligands, myostatin (GDF-8) and Activin A, are potent negative regulators of skeletal muscle growth^[14]^. These ligands bind to ActRIIA/B, triggering the phosphorylation of SMAD2/3, which subsequently inhibits the Akt/mTOR protein synthesis pathway and activates the ubiquitin-proteasome proteolytic system^[15]^. Genetic deletion or pharmacological inhibition of ActRII receptors has consistently demonstrated significant muscle hypertrophy and improved metabolic profiles in preclinical models^[16–18]^.

Activins are dimeric proteins, mainly including Activin A and Activin B, and exert multiple physiological functions by binding to ActRII. In skeletal muscle, Activin A signals through ActRII, inhibiting muscle cell differentiation and promoting protein catabolism, thereby acting as a negative regulator of skeletal muscle mass. Together with myostatin, it constitutes a key signal inhibiting muscle growth. Studies have shown that combined blockade of GDF8 and Activin A achieves muscle growth effects comparable to direct blockade of ActRIIA/B receptors, suggesting that Activin A is the second most important negative regulator of muscle that acts synergistically with GDF8. In adipose tissue, activin promotes lipid storage via ActRII, and blocking this signaling pathway promotes fat metabolism^[19]^. In adipocytes, Activin A is one of the key signaling molecules that promote lipid storage. Blocking its binding to ActRII receptors exerts two major beneficial effects: on one hand, it directly relieves the inhibitory effect of Activin A on adipocyte differentiation, allowing more preadipocytes to differentiate and mature normally, thereby optimizing adipose tissue function; on the other hand, it relieves the inhibition of lipolysis by Activin A, promoting the breakdown of triglycerides into free fatty acids and glycerol for consumption, directly reducing fat accumulation from the source and increasing its expenditure.

Myostatin (GDF8) is the most well-studied negative regulator of muscle in the TGF-β superfamily, acting through ActRIIA/B receptors. Myostatin is mainly expressed by skeletal muscle cells and negatively regulates myoblast growth. Myostatin binds with high affinity to the ActRIIB receptor and initiates signaling through the Smad2/3-dependent pathway. Myostatin knockout mice exhibit a significant increase in muscle mass due to muscle hyperplasia and hypertrophy^[20]^. In mammals, loss-of-function mutations in the MSTN gene are closely associated with increased skeletal muscle mass in cattle, sheep, dogs, and humans. Elevated myostatin levels are thought to be associated with various muscle-wasting conditions such as disuse atrophy, AIDS-related cachexia, cancer-related cachexia, and age-related sarcopenia. In adipose tissue, myostatin has also been shown to regulate adipogenesis by directly or indirectly blocking BMP signaling. The most prominent feature of myostatin is that its primary action is largely restricted to skeletal muscle, with relatively minor systemic effects, making it an ideal target for muscle-gain strategies.

Growth differentiation factor 11 (GDF11) is a secreted factor of the TGF-β family and is evolutionarily the closest relative to myostatin. Both have similar binding affinities for the ActRIIB receptor, suggesting they should have comparable mechanisms of action at the signal transduction level. However, research on the function of GDF11 is controversial. Early high-impact studies suggested that GDF11 levels decline with age and that supplementing GDF11 to “young” levels is beneficial for various age-related diseases, particularly showing an effect opposite to that of myostatin on skeletal muscle regeneration^[21]^. However, most independent laboratories have not been able to replicate the finding that GDF11 levels decline with age in subsequent studies. All follow-up studies have described that GDF11 inhibits skeletal muscle regeneration, and that high-dose GDF11 can induce skeletal muscle atrophy and cachexia. In addition, several studies^[22]^ have examined the effects of GDF11 and/or its downstream ActRII pathway on cardiac function and bone biology. GDF11 is a factor with potential cardiovascular regulatory and bone-protective effects, and enhanced blockade of GDF11 and/or its downstream ActRII pathway may produce further effects on skeletal development and cardiovascular regulation.

Bone morphogenetic proteins (BMPs) are a diverse group of members of the TGF-β superfamily. ActRII also binds a subset of BMPs, including BMP2, BMP4, BMP7 (also known as OP-1), BMP9, and BMP10^[23]^. BMP signaling plays indispensable regulatory roles in various physiological processes including bone formation, embryonic development, hematopoiesis, and iron metabolism. The ligand diversity and functional complexity of ActRII dictate that drug development targeting this node requires careful consideration of the balance among multiple aspects: “which ligands to block,” “which receptor subtype(s) to block,” and “which physiological functions to preserve.” Research to date indicates that GDF8 and Activin A are the two primary negative regulators of muscle mass; BMPs can influence muscle via the ActRII pathway while also playing important roles in bone formation; understanding the functions of different ActRII ligands and their interactions is of great importance for designing anti-ActRII antibodies with an optimal risk-benefit profile.

Previous clinical efforts, most notably with bimagrumab (a first-generation anti-ActRII antibody), have validated this mechanism. In patients with type 2 diabetes and obesity, bimagrumab treatment led to significant fat loss and concomitant lean mass gain^[24]^. The BELIEVE phase 2 trial demonstrated that the combination of bimagrumab and semaglutide significantly enhances weight loss compared to monotherapies while potentially improving body composition in adults with obesity^[25]^. However, earlier iterations of ActRII-targeted therapies have faced challenges, including suboptimal potency, limited differentiation in “real-world” combination settings, and developability hurdles such as high surface hydrophobicity, which increases the risk of self-aggregation and complicates high-concentration subcutaneous formulation. Furthermore, as the industry moves toward combination therapies, there is a critical need for next-generation antibodies with superior binding affinity and optimized physicochemical properties to support patient-friendly dosing regimens.

Recent advances in AI-driven antibody design—encompassing diffusion models, language models, and structure-guided generation—have achieved remarkable progress in affinity maturation, pushing binding constants from nanomolar to picomolar levels across platforms such as RFantibody^[26]^, Chai-2^[27]^, Jam-2^[28]^ and Boltzgen^[29]^. The RFdiffusion framework, for instance, demonstrated atomically accurate *de novo* design of VHHs and full antibodies against user-specified epitopes, while Chai-2 achieved double-digit hit rates in fully de novo antibody design across 52 diverse targets, and JAM-2 generated drug-like antibodies with high success rates across 16 unseen targets. These and other efforts reflect a growing shift from “discovery” to “design”, raising the possibility that the long-standing bottleneck of generating high-affinity binders may soon be overcome.

However, as high-affinity designs have proliferated, a more subtle and clinically relevant bottleneck has emerged: affinity alone is a poor predictor of functional activity. A 2023 Nature study directly compared antibodies against the same epitope on CD40 with K_D_ values ranging from 5.22 nM to 925 nM, and found that medium affinity (∼50 nM) induced the strongest immune activation, while higher affinity paradoxically reduced function a strategy successfully extended to 4-1BB and PD-1^[30]^. Similar observations have been made in T-cell receptor systems, where antigen response follows an inverted U-shaped “bell curve”: excessively high affinity can trap receptors in non-signaling states or trigger rapid internalization and negative feedback^[31]^. Beyond T-cell biology, recent functional studies on GIPR antibodies have shown that increasing binding affinity by 7–10-fold does not translate into proportional gains in antagonistic potency, with IC₅₀ values remaining essentially unchanged across multiple affinity-enhanced variants^[32]^. These examples underscore that the ability to bind tightly — even with atomic precision — is necessary but far from sufficient. What ultimately matters are whether an antibody can effectively modulate cellular signaling *in situ*, a property shaped by epitope accessibility, binding geometry, stability under physiological conditions, and the nuanced architecture of the target membrane protein. The emerging literature thus calls for a fundamental reorientation in the evaluation of AI-designed antibodies: from isolated affinity metrics (K_D_) toward integrated functional readouts that better reflect *in vivo* activity. This shift is not merely semantic — it has direct engineering consequences. An antibody may achieve picomolar affinity in SPR, yet fail to block ligand-induced signaling in cell-based assays — either because it binds an epitope that does not disrupt receptor activation, or because its instability under physiological conditions compromises its functional activity.

Recognizing this “high-affinity trap”, we have developed AlfaDAX, integrated AI-driven platforms designed to accelerate antibody discovery and optimization from a function-first perspective. Unlike conventional workflows that prioritize affinity screening as the primary filter, AlfaDAX incorporate multi-objective optimization that simultaneously evaluates predicted binding energy, epitope accessibility, and cellular functional activity. In the present study, we apply the AlfaDAX platform to a different target, ActRIIA/B, with a distinct objective: rather than pursuing further affinity gains, we aim to enhance cellular functional activity on an identical epitope, using point mutations guided by AI-predicted functional and energy metrics, without generating novel antibody sequences. Our goal is to improve functional blocking activity while maintaining favorable developability parameters. Applying this strategy, we identified AB130-165, a differentiated high-affinity humanized monoclonal antibody that exhibits 9.5-fold superior functional blockade of Activin A/ActRII signaling compared to the bimagrumab, along with lower non-specific binding. In diet-induced obese (DIO) mice, AB130-165 monotherapy, and more significantly, its combination with semaglutide, resulted in profound body composition improvements. The combination therapy not only accelerated fat loss but also preserved lean mass/body weight percentage to a degree significantly greater than that of the bimagrumab combination. These results provide a robust scientific rationale for AB130-165 as a potential Best-in-Class candidate for achieving high-quality weight loss and addressing the unmet medical need for muscle-preserving obesity therapies. Our findings demonstrate that dual ActRIIA/B blockade synergizes with GLP-1RA treatment to enhance fat loss while preserving lean mass, supporting a differentiated approach for next-generation obesity therapies focused on improving body composition.

## 2 Results

### 2.1 Optimization of Cellular Function of Anti-ActRIIA/B Antibodies via the AlfaDAX Platform

To break the traditional engineering trade-offs between antibody potency and developability, we employed the AlfaDAX *in silico* optimization platform to enhance the functional blocking activity of anti-ActRIIA/B antibodies without altering their target epitopes significantly. Starting from a parental humanized clone (bimagrumab) with established dual-binding capabilities to ActRIIA and ActRIIB, we performed three iterative rounds of structure-guided point mutation design. This workflow was driven by AI-predicted functional and energy metrics to enrich variants with superior bioactivity while concurrently filtering out potential developability liabilities (e.g., aggregation risk, non-specific binding). (Figure 1A). Candidate variants selected from each *in silico* screening round were expressed in CHO-K1 cells and subjected to wet-lab validation, focusing on cellular functional potency (IC₅₀) in an *ACVR2A/B*-overexpressing SMAD-responsive luciferase reporter cell line. As shown in Figure 1B, the functional blocking activity improved progressively across the three rounds. The lead candidate from Round 3, designated AB130-165, exhibited an IC₅₀ approximately 10-fold lower than that of the parental bimagrumab analog, representing a substantial gain in cellular function on the same epitope. Structural modeling of the antibody–receptor complexes (Figure 1C) further suggested that the AlfaDAX-introduced mutations optimized the functional interface and spatial positioning without shifting the core binding epitope as evidenced by the preserved epitope footprint compared to the published crystal structure of bimagrumab (PDB: 5NH3). Importantly, the AI-predicted binding free energy (ΔG) for AB130-165 was dramatically lower than that of the parental antibody. As summarized in Table 1, AB130-165 achieved an energy score of –16.81 kcal/mol for ActRIIA and –16.39 kcal/mol for ActRIIB, compared to –10.70 kcal/mol and –14.52 kcal/mol for bimagrumab, respectively. Notably, another candidate from the same optimization round, AB130-188, exhibited even more favorable predicted binding energies: −18.48 kcal/mol for ActRIIA and −18.03 kcal/mol for ActRIIB, suggesting an even stronger predicted interaction. All candidates maintained high abPTM scores (≥0.90) and identical block_rate scores (0.91), confirming that the optimized antibodies bind the same functional epitope and are predicted to effectively block ligand engagement. To experimentally validate this prediction, we measured the equilibrium dissociation constants (Kᴅ) by surface plasmon resonance (SPR) using a Biacore T200 instrument (Figure 5). Remarkably, AB130-165 exhibited a Kᴅ of 0.204 pM for ActRIIA – a 1,054-fold improvement over bimagrumab (215 pM) and a Kᴅ of 0.243 pM for ActRIIB, which was 10-fold higher than that of bimagrumab (2.55 pM). Consistent with its more favorable predicted ΔG, AB130-188 demonstrated even higher affinity, with Kᴅ values of 0.00849 pM for ActRIIA and 0.00108 pM for ActRIIB, representing an additional 24-fold and 225-fold improvement over AB130-165, respectively These SPR data directly link the AI-predicted energy lowering to experimentally confirmed ultra-high affinity, demonstrating that AlfaDAX successfully drove both functional and affinity maturation. Notably, despite the dramatic affinity improvement (up to ∼1,000-fold for AB130-165 and even higher for AB130-188), the cellular functional potency (IC₅₀) increased only ∼10-fold, not proportionally with affinity. This observation indicates that beyond a certain threshold, further increases in affinity yield diminishing returns in cellular blockade – a classic example of the “high-affinity trap”. For AB130-188, despite its ∼25-fold higher affinity compared to AB130-165, the functional IC₅₀ was even weaker than AB130-165 (Figure 2), further supporting the notion that once receptor occupancy is saturated, additional binding strength does not translate into enhanced signaling blockade. Moreover, the antibody’s developability parameters (Supplementary Figure 1) remained favorable, with no increase in aggregation risk or non-specific binding. Together, these findings validate the AlfaDAX-enabled “functional maturation on the same epitope” strategy, successfully bypassing the naïve pursuit of affinity alone and yielding AB130-165 – a candidate with both ultra-high affinity (sub-picomolar) and significantly enhanced cellular function.

**Figure 1.**
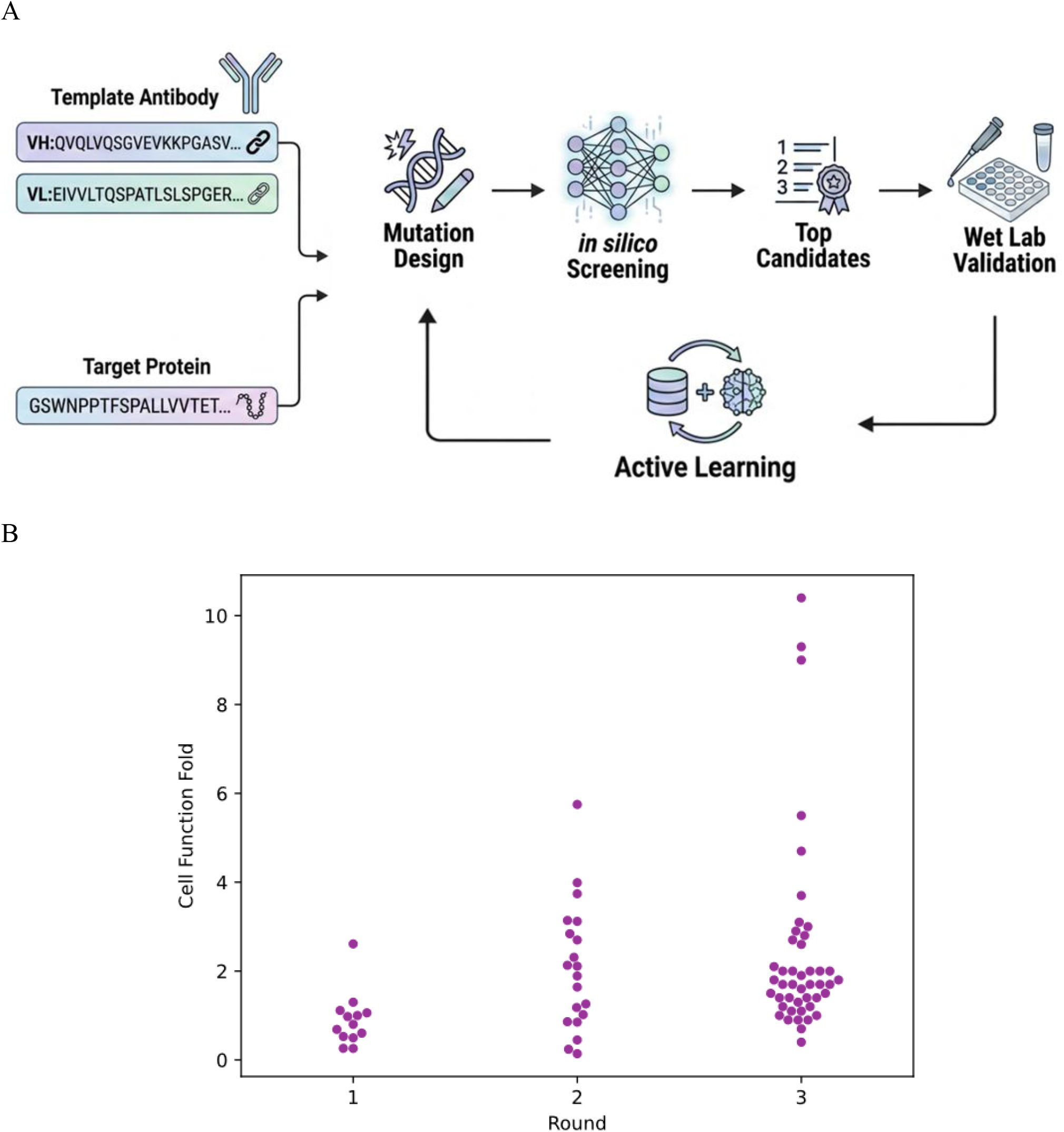
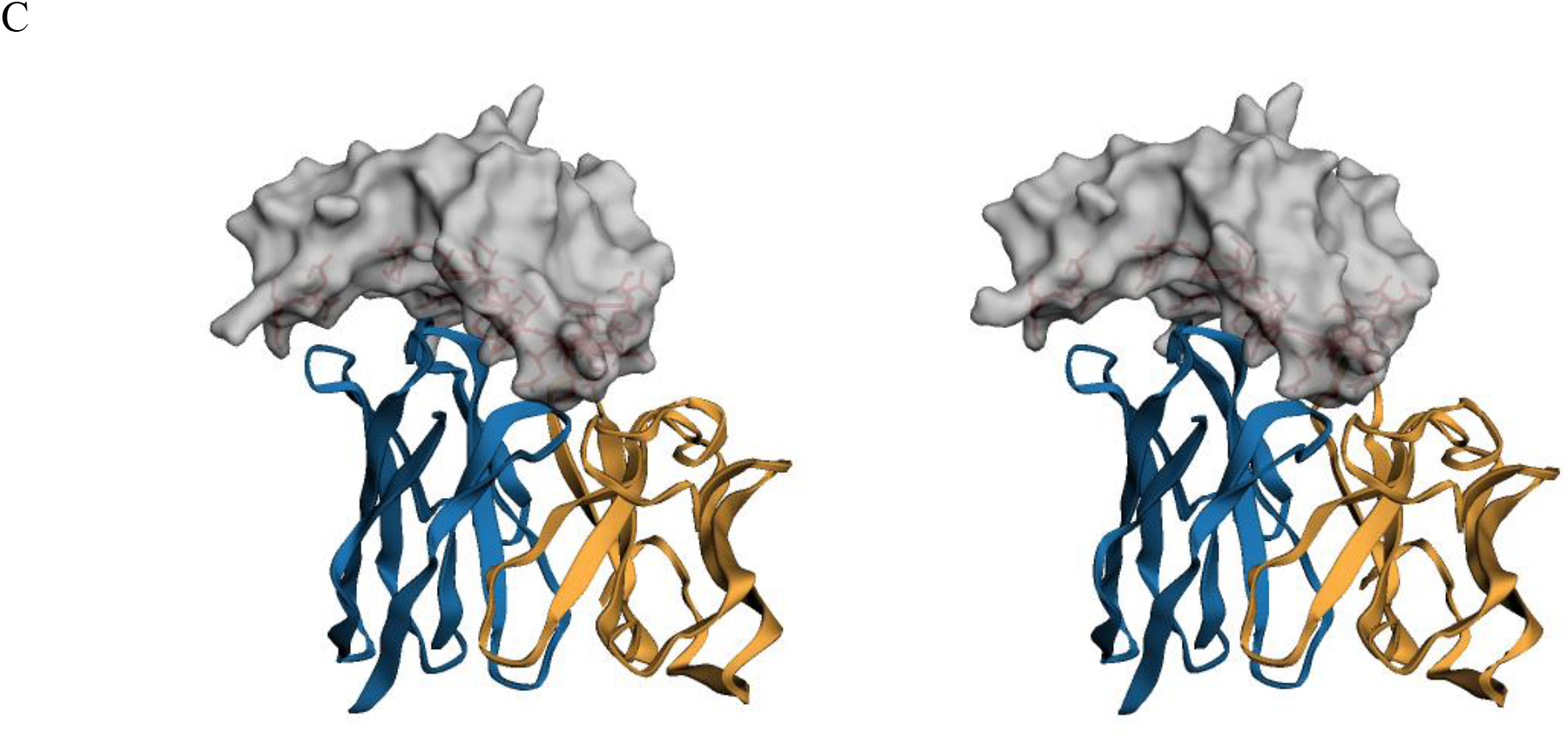
Design and *in silico* optimization of anti-ActRIIA/B antibodies via the AlfaDAX platform. (A) Schematic workflow of the AlfaDAX iterative *in silico* optimization platform. Starting from a parental anti-ActRIIA/B antibody, three rounds of point mutation design were performed, guided by AI-predicted DAX functional scores. Selected candidates were expressed and subjected to *in vitro* functional validation. (B) Cellular functional activity of designed antibodies across rounds 1 to 3, quantified as IC₅₀ in an ActRIIA/B reporter cell line. IC₅₀ values decreased progressively, representing an approximately 10-fold improvement in functional blocking activity from round 1 to round 3. (C) Structural prediction of antibody–ActRIIA complexes. Left: Predicted binding mode of AB130-165 in complex with ActRIIA. Blue: heavy chain; orange: light chain. Gray surface: ActRIIA; red sticks: key epitope residues. Right: Crystal structure of bimagrumab in complex with ActRIIA (PDB: 5NH3) for comparison. The designed antibody AB130-165 binds to a similar epitope on ActRIIA as the reference antibody bimagrumab, despite sequence differences in CDR regions.

**Figure 2.**
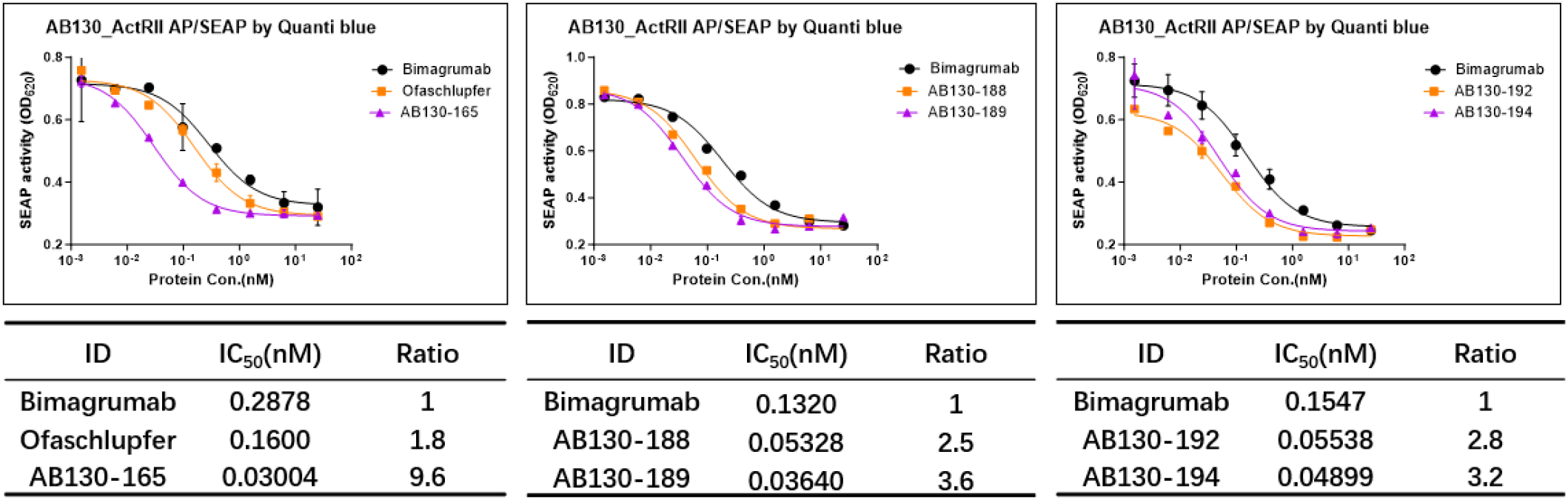
Blocking activity of candidate antibodies against activin A-induced SMAD signaling in HEK-Blue™ TGF-β reporter cells overexpressing ActRIIA and ActRIIB. Secreted alkaline phosphatase (SEAP) activity in the cell supernatant was measured using Quanti Blue reagent, expressed as OD_620_. Cells were co-treated with serial dilutions of candidate antibodies and activin A, and dose-response curves were generated. Half-maximal inhibitory concentrations (IC₅₀) were calculated by four-parameter logistic fitting. The ratio indicates the fold improvement in IC₅₀ relative to bimagrumab (set to 1).

**Figure 3.**
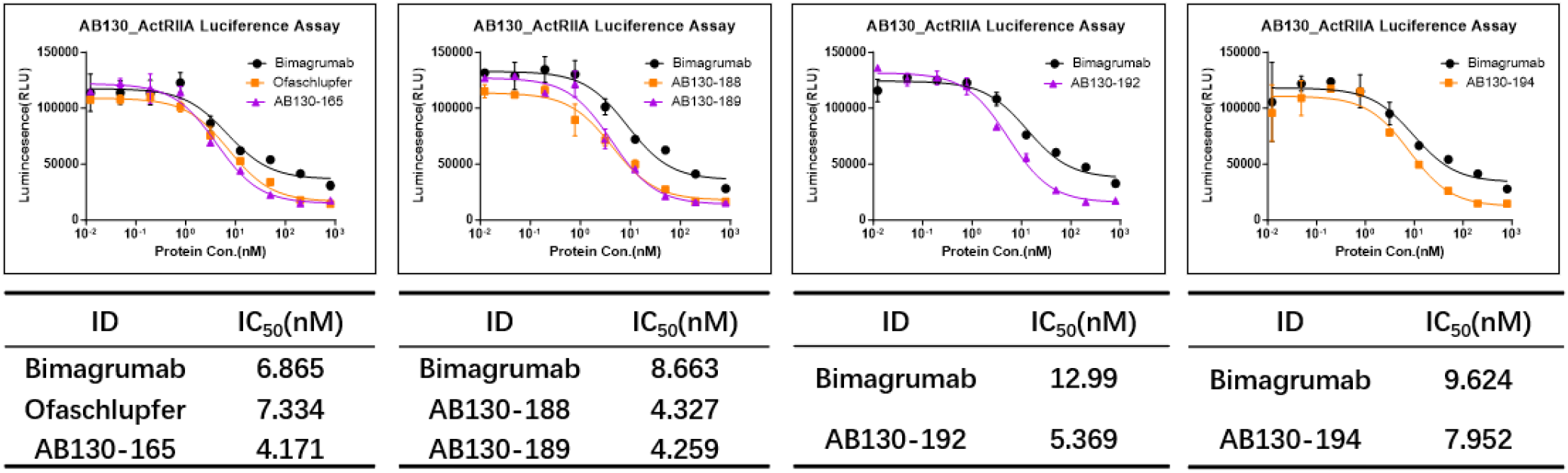
Blocking activity of candidate antibodies against activin A-induced SMAD signaling in an ActRIIA-specific reporter cell line. The H_ActRIIA reporter cell line (*ACVR2B*-knockout, *ACVR2A*-overexpressing) from Genomeditech was used to evaluate the blocking effect of candidate antibodies on the ActRIIA/activin A signaling axis. Cells were co-treated with serial dilutions of candidate antibodies and activin A. SEAP activity in the cells was measured using Luciference, and dose-response curves were generated to calculate half-maximal inhibitory concentrations (IC₅₀). Data are presented as mean ± SEM.

**Figure 4.**
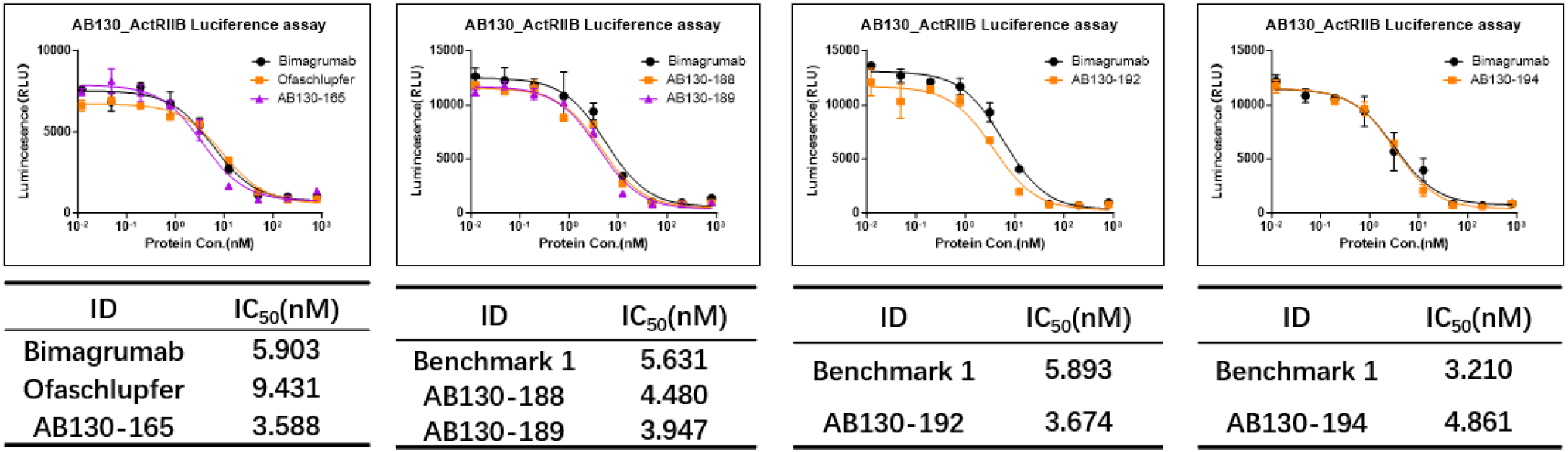
Blocking of activin A-induced SMAD signaling by candidate antibodies in an ActRIIB-specific reporter cell line. An *ACVR2A*-knockout, *ACVR2B*-overexpressing H_ActRIIB reporter cell line (Genomeditech) was used. Cells were treated with increasing concentrations of candidate antibodies together with activin A. SEAP activity was measured via luciferase, and IC₅₀ values were derived from dose-response curves. Results are presented as mean ± SEM

**Figure 5.**
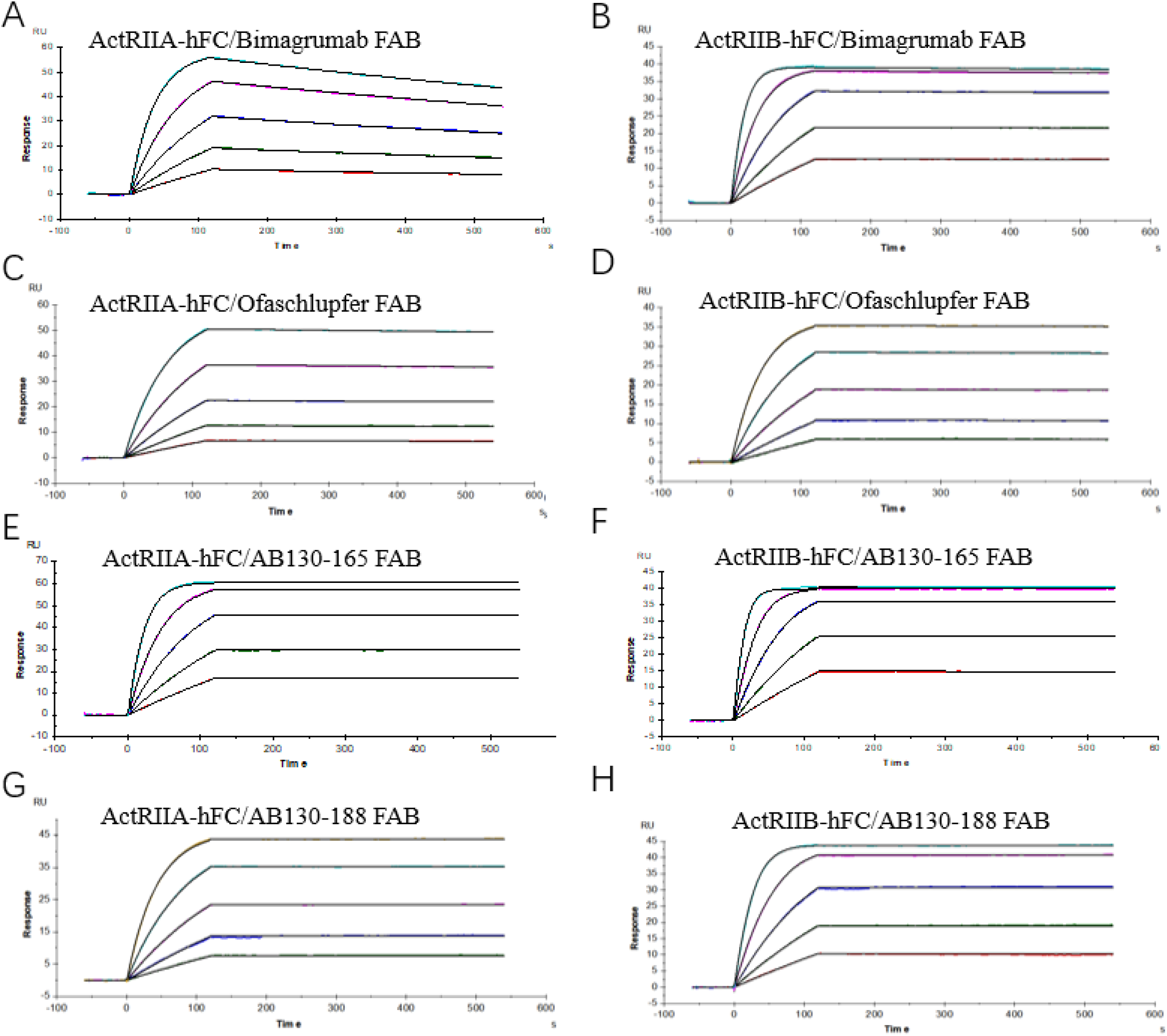
Characterization of the affinity of AB130-165/188 and control molecules. (A). SPR determination of the affinity of bimagrumab for ActRIIA. The ActRIIA-hFc target protein was immobilized on a Protein A chip, and the Fab antibody was tested at a starting concentration of 10 nM with five serial 2-fold dilutions and a blank concentration. The remaining samples were tested using a similar method. (B). SPR determination of the affinity of bimagrumab for ActRIIB. For the ActRIIB affinity measurement, the starting antibody concentration was 5 nM, with five serial 2-fold dilutions and a blank concentration. (C) SPR determination of the affinity of Ofaschlupfer for ActRIIA. (D) SPR determination of the affinity of Ofaschlupfer for ActRIIB. (E). SPR determination of the affinity of AB130-165 for ActRIIA. (F). SPR determination of the affinity of AB130-165 for ActRIIB. (G). SPR determination of the affinity of AB130-188 for ActRIIA. (H). SPR determination of the affinity of AB130-188 for ActRIIB.

**Table 1.**
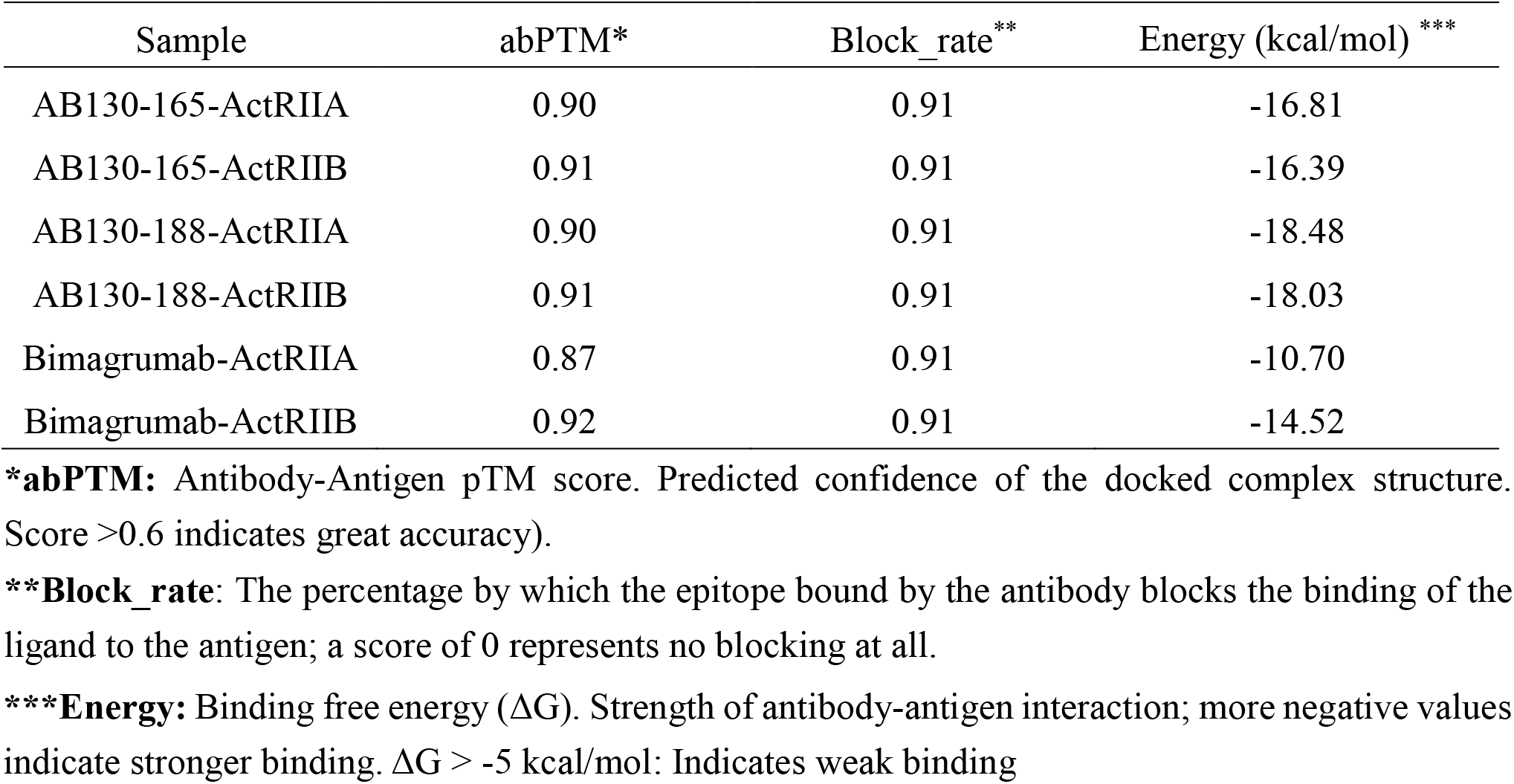
Confidence metrics of AB130-165/188 and bimagrumab.

### 2.2 *In vitro* functional validation of candidate molecules

After three rounds of optimization and screening using the AlfaDAX platforms (data not shown in detail), five optimal molecules were selected for comprehensive evaluation. During the evaluation, bimagrumab (laboratory-prepared, with a sequence identical to GSRS Record UNII N15SW1DIV8) and Ofaschlupfer (Sixpeaks, patent number WO2025027052A1) were used as positive controls and prepared following the same manufacturing process.

To evaluate the blocking effect of the candidate antibodies on the SMAD signaling pathway, ActRIIA and ActRIIB were overexpressed in HEK-Blue™ TGF-β reporter cells. Following functional evaluation, the IC₅₀ values of the five candidate molecules for blocking the activin A signaling pathway were 2.5- to 9.6-fold lower than that of bimagrumab, indicating a corresponding improvement in potency. Among them, AB130-165 emerged as the most potent candidate, with an IC₅₀ of 0.03004 nM, representing a 9.6-fold improvement in blocking potency compared with bimagrumab (0.2878 nM) (Figure 2). To further dissect the blocking effects specifically mediated by ActRIIA or ActRIIB, we used H_ActRIIA Reporter Cell Line and H_ActRIIB Reporter Cell Line from Genomeditech. In these two cell lines, either ActRIIA or ActRIIB was knocked out, and the other receptor was overexpressed, thereby generating SMAD signaling reporter cells with selective receptor expression. Using the H_ACTRIIA reporter cell line (which overexpresses ActRIIA and lacks ActRIIB), we confirmed that the candidate molecules primarily enhanced the blockade of the ActRIIA/activin A signaling axis. The IC₅₀ of AB130-165 was 4.17 nM, compared with 7.36 nM for Ofaschlupfer and 6.92 nM for bimagrumab (measured as 6.92 nM on the same 96-well plate; the mean IC₅₀ of bimagrumab across four plates was 9.69 nM). In addition, bimagrumab showed a lower maximum blocking rate at the highest concentration. The remaining four candidate molecules exhibited similar results (Figure 3). Concurrently, we used the H_ActRIIB reporter cell line (which overexpresses ActRIIB and lacks ActRIIA) to evaluate the blocking activity against the ActRIIB/activin A signaling pathway. The results showed that the IC₅₀ values of the two control molecules and the five candidate molecules were essentially comparable, with differences within twofold, and the maximum blocking rates were also similar (Figure 4). Thus, through optimization using the AlfaDAX platform targeting the ActRIIA/activin A signaling blockade, we obtained molecules that were up to 9.6-fold superior to the control at the molecular level.

The candidate molecule AB130-165/188 was selected and submitted to a third-party laboratory for surface plasmon resonance (SPR) affinity measurement using a Biacore T200 instrument. The results showed that AB130-165 exhibited an affinity of 0.204 pM for ActRIIA, representing a 1054-fold improvement compared to bimagrumab. For ActRIIB, the affinity was 0.243 pM, which was 10-fold higher than that of bimagrumab (Figure 5). Subsequently, the five most promising candidate molecules were selected for further validation of their binding activity at the molecular level. The half-maximal effective concentration (EC_50_) of the candidate molecules for binding to the target proteins was determined by ELISA. The data indicated that the EC_50_ values of all molecules were essentially similar, with differences within twofold. In addition, cell lines overexpressing ActRIIA and ActRIIB were established, and the EC_50_ of the candidate antibody molecules binding to the cell-surface target proteins was measured. The results again showed no significant differences, as presented in Table 2.

**Table 2.** Binding Activity Statistics of Candidate Molecules.

| Sample ID | ActRIIA |  |  |  | ActRIIB |  |
| --- | --- | --- | --- | --- | --- | --- |
|  | K<sub>D</sub> (pM) | ELISA EC<sub>50</sub>(nM) | Cell Binding EC<sub>50</sub>(nM) | K<sub>D</sub> (pM) | ELISA EC<sub>50</sub> (nM) | Cell Binding EC<sub>50</sub> (nM) |
| Bimagrumab | 215 | 0.1124 | 2.722 | 2.55 | 0.1542 | 0.6106 |
| Ofaschlupfer | 34.6 | 0.1090 | 1.676 | 3.19 | 0.1106 | 0.4403 |
| AB130-165 | 0.204 | 0.1220 | 3.217 | 0.243 | 0.1430 | 0.6172 |
| AB130-188 | 0.00849 | 0.1335 | 2.142 | 0.00108 | 0.1191 | 0.4998 |
| AB130-189 | N/A* | 0.1504 | 1.766 | N/A* | 0.1863 | 0.4864 |
| AB130-192 | N/A* | 0.1109 | 2.347 | N/A* | 0.1431 | 0.3504 |
| AB130-194 | N/A* | 0.1795 | 1.719 | N/A* | 0.1535 | 0.3920 |

\*: Not determined.

While optimizing antibodies, we simultaneously analyzed whether the developability of the recommended molecules fell within normal ranges, and we performed a series of developability assessments. The expressibility of the antibodies was evaluated using CHO-K1 cells, and the results showed that all candidate molecules exhibited expression levels within a normal range (expression >200 mg/L at day 4). The aggregation stability of the candidate antibody molecules was analyzed by SEC-HPLC, and the results indicated that the purity of all molecules was above 95%. Hydrophobicity of the antibodies was analyzed by HIC (hydrophobic interaction chromatography), and the candidate antibodies were comparable to bimagrumab. Finally, we measured the Tm and Tagg values of the antibodies to assess their thermal stability, and the results showed that all candidate molecules had values within the normal range (>65°C) (Table 3). These results demonstrate that our DAX platform can predict molecule developability while recommending optimized molecules, thereby achieving simultaneous optimization of both molecular activity and developability.

**Table 3.** Developability analysis of candidate molecules.

| Sample ID | Transient Expression in CHO Cells (mg/L) | SEC-HPLC (%) | HIC Retention Time (min) | Tm (°C) | Tagg (°C) | HEK_293T Non-specific Binding MFI |
| --- | --- | --- | --- | --- | --- | --- |
| Bimagrumab | 333.8 | 97.09% | 20.3 | 72.60 | 70.27 | 1208 |
| AB130-165 | 312.0 | 97.17% | 20.7 | 69.30 | 66.45 | 813 |
| AB130-188 | 291.5 | 97.80% | 20.7 | 67.88 | 66.02 | 726 |
| AB130-189 | 207.0 | 97.96% | 20.2 | 65.20 | 65.67 | 925 |
| AB130-192 | 336.5 | 98.08% | 19.8 | 68.10 | 66.35 | 845 |
| AB130-194 | 321.0 | 98.07% | 20.3 | 67.70 | 66.34 | 620 |

### 2.4 ActRII modulation improves weight-loss quality in DIO mice

Human and mouse ActRII proteins are highly conserved, sharing an extremely high degree of sequence identity, and semaglutide also retains full activity in mice, diet-induced obese mice were generated by feeding with a 60% high-fat diet for 16-19 weeks. Mice were treated for 27 days (Figure 6A) with subcutaneous injections of semaglutide (0.123 mg/kg) or vehicle control, ActRII blocking antibodies (20 mg/kg) or the combination of semaglutide and ActRII antibodies (In this manuscript, only the results for AB130-165 were presented). Body weight and food consumption were measured twice a week, and body composition was assessed by NMR at days 0, 13, 20 and 27. Body weight changes were expressed as percentage increase from baseline. Over the 4-week treatment period, the NC group showed a weight gain of 5.5%, whereas the AB130-165 and bimagrumab groups exhibited increases of 7.5% and 11.0%, respectively. Semaglutide monotherapy reduced body weight by 24.3% compared to baseline, while the addition of AB130-165 or bimagrumab further increased weight loss to 33.4% and 25.5%, respectively (Figure 6B). The cumulative food intake in each group was consistent with the observed body weight changes (Figure 6C). Body composition changes were analyzed to assess whether selective ActRII blockade improves the quality of semaglutide-induced weight loss (Figure 6D, 6E). Critically, semaglutide alone led to substantial muscle loss (−15.1%) and a 46.3% fat reduction. Combination with AB130-165 markedly attenuated muscle loss (−9.3%) and dramatically enhanced fat reduction (−77.8%). In contrast, the bimagrumab combination resulted in less muscle loss (−6.3%) but also less fat reduction (−65.0%). The superior quality of weight loss achieved by AB130-165 plus semaglutide was further reflected in body composition (Table 4), yielding 67.4% lean mass and 12.6% fat, compared to semaglutide alone (54.3% lean, 26.1% fat) and the bimagrumab plus semaglutide (62.3% lean, 17.4% fat). Taken together, these findings demonstrate that both ActRIIA and ActRIIB blockade with antibody AB130-165 improves weight-loss quality by preserving lean mass and maximizing fat reduction, outperforming both semaglutide alone and the combination with bimagrumab.

**Figure 6.**
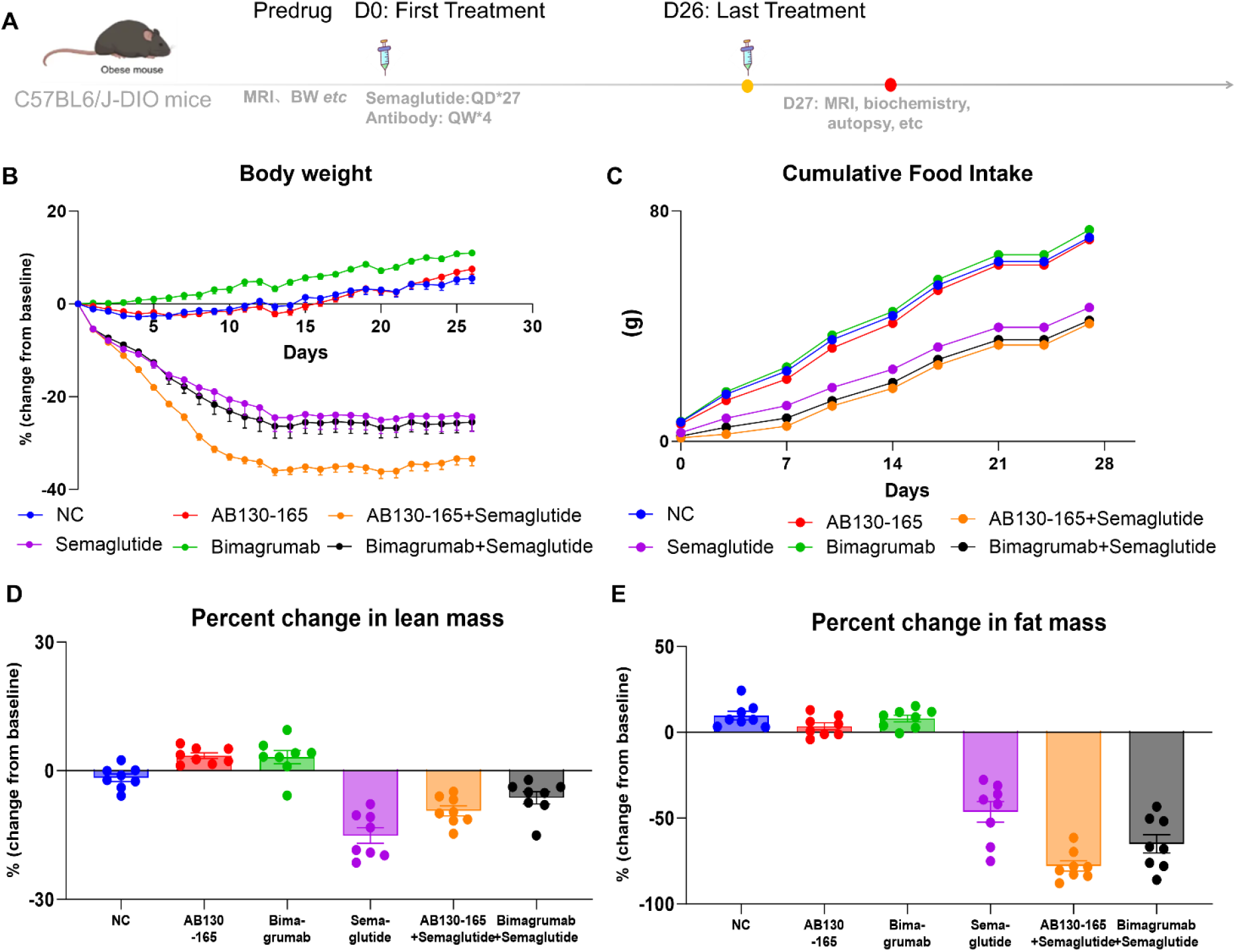
Both ActRIIA and ActRIIB blockade improves weight-loss quality under semaglutide treatment in DIO mice. (A) Experimental schematic. (B) Percent change in total body weight from baseline over 27 days. (C) Cumulative food intake. (D) Percent change in lean mass. (E) Percent change in fat mass. Data are presented as mean ±SEM (n = 8 per group).

**Table 4.** Day 27 Body Composition Phenotype (DIO Mice)

| Group | Lean Mass/BW (%) | Fat Mass/BW (%) |
| --- | --- | --- |
| Semaglutide Monotherapy | 54.4 | 26.1 |
| AB130-165 + Semaglutide | 67.4 | 12.6 |
| Bimagrumab + Semaglutide | 62.3 | 17.4 |

To evaluate the effects of ActRII blockade and semaglutide on white adipose tissue and skeletal muscle, we dissected and weighed the inguinal, epididymal, and mesenteric fat depots, as well as the gastrocnemius, tibialis anterior, and quadriceps muscles at the end of the treatment period. AB130-165 in combination with semaglutide induced the greatest fat reduction across all three depots examined. Compared with semaglutide monotherapy, the combination of AB130-165 with semaglutide resulted in additional reductions of 35.1% in inguinal fat weight, 33.1% in epididymal fat weight, and 19.2% in mesenteric fat weight; the corresponding values for the bimagrumab plus semaglutide group were 21.7%, 15.2%, and 0.6%, respectively (Figure 7). AB130-165 alone and in combination with semaglutide maintained muscle mass comparable to that observed with bimagrumab, whereas semaglutide monotherapy significantly reduced muscle mass. Taken together, AB130-165 plus semaglutide uniquely achieved maximal fat loss while preserving lean mass, supporting AB130-165 as a superior combination partner with semaglutide for the dual goal of fat reduction and muscle preservation.

**Figure 7.**
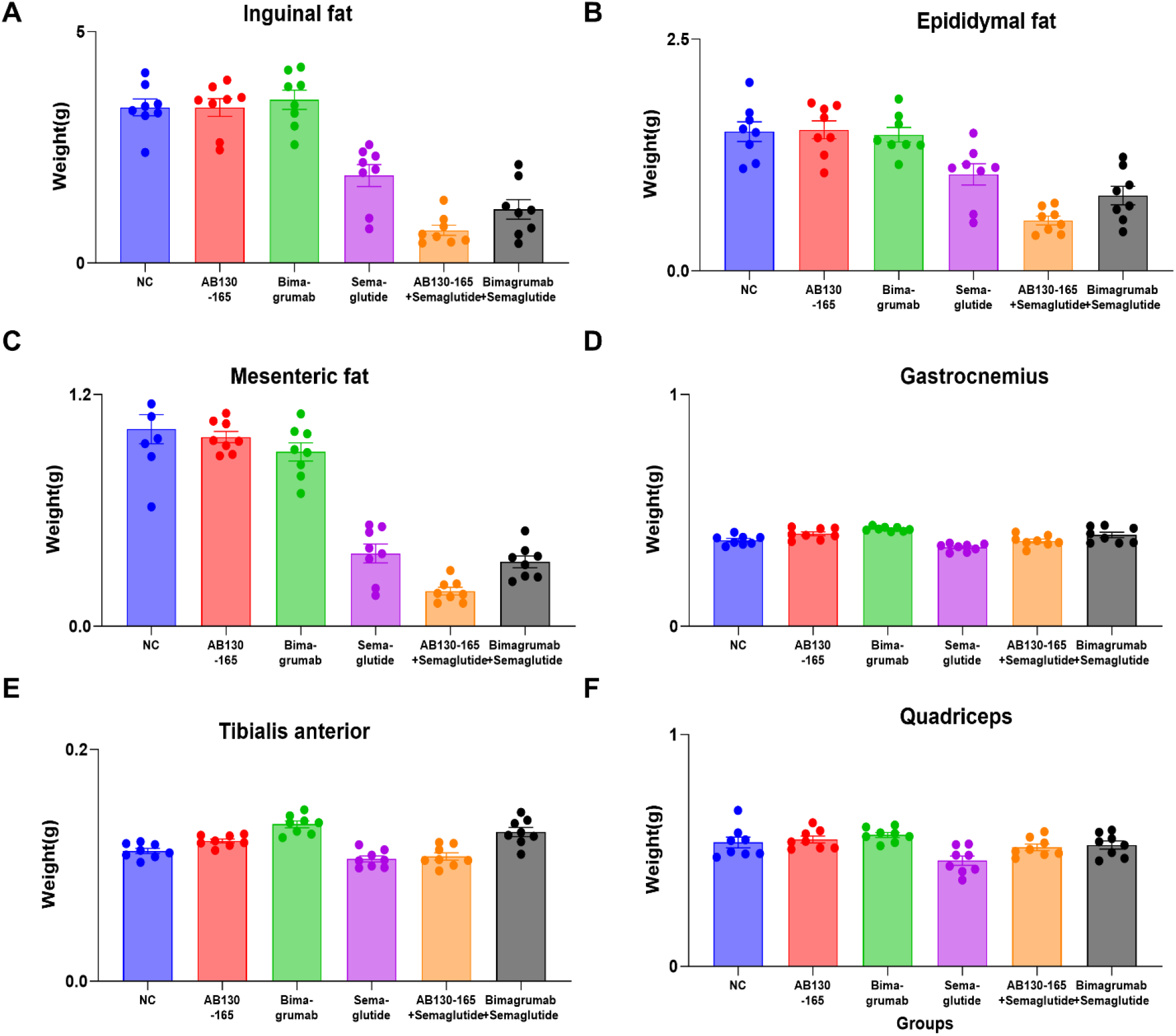
ActRII inhibition combined with semaglutide leads to a further reduction in adipose tissue mass and preserves muscle mass during weight loss. Fat depot weights of each group. (A) inguinal, (B) epididymal, (C) mesenteric fat depots (D) gastrocnemius, (E)tibialis anterior, and (F) quadriceps muscles. Data are presented as mean ±SEM (n = 8 per group).

Serum lipid profiles were analyzed to assess the metabolic effects of the interventions. AB130-165 alone reduced triglycerides by 42% and LDL-C by 17%, outperforming bimagrumab (TG: –16%; LDL-C: –11%) (Figure 8). Consistent with previous reports, semaglutide monotherapy significantly improved the lipid profile, leading to substantial reductions in TC (39%), TG (28%), and LDL-C (49%). Combination therapies further amplified these beneficial effects. Notably, AB130-165 plus semaglutide exhibited greater efficacy in ameliorating TC (–42% vs. –38%) and LDL-C (–59% vs. – 51%) compared to the bimagrumab plus semaglutide regimen. In contrast, all treatments, particularly those containing semaglutide, resulted in decreased HDL-C levels (AB130-165 plus semaglutide: – 33%; bimagrumab plus semaglutide: –31%), with comparable effects between the two combinations. Thus, AB130-165 combined with semaglutide confers superior cardiometabolic benefits over the bimagrumab-based regimen.

**Figure 8.**
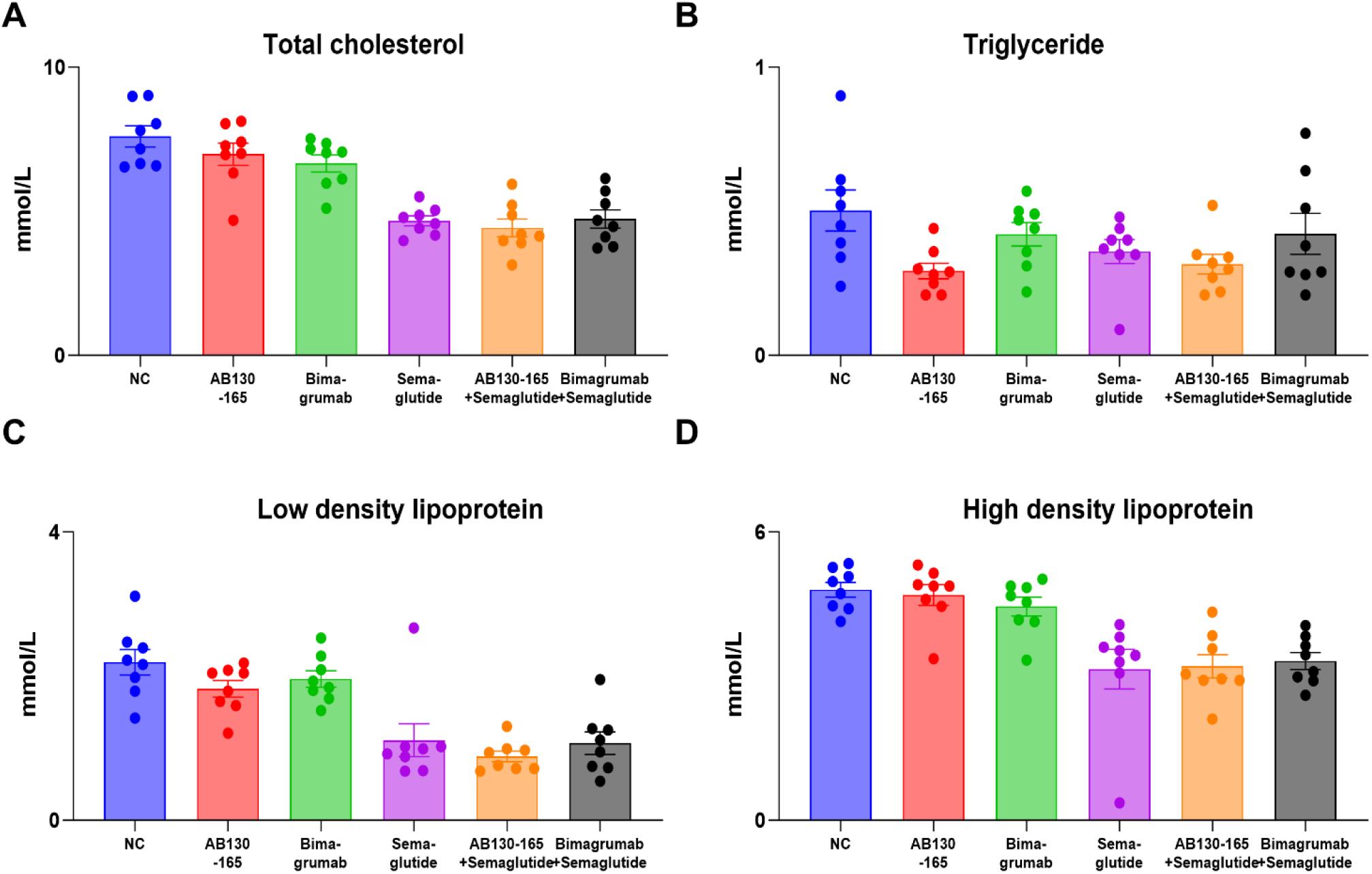
Combined ActRII blockade (AB130-165) plus GLP-1 receptor agonism (semaglutide) improves circulating lipid profiles beyond GLP-1 monotherapy. (A) total cholesterol, (B) Triglyceride, (C) Low density lipoprotein (D) High density lipoprotein. Data are presented as mean ± SEM (n = 8 per group).

## 3 Methods

### 3.1 Protein Expression

The AI-recommended amino acid sequence was optimized for the codon preference of *Cricetulus griseus* (CHO). The heavy and light chains of the antibody were separately constructed into the pCDNA3.4 vector through gene synthesis. The heavy and light chain plasmids were co-transfected into CHO-K1 cells for cultivation and expression. The supernatant of the expressed cells was harvested and purified using Protein A purification resin (Cytiva).

### 3.2 Affinity determination of candidate molecules

Antibody-antigen binding kinetics were assessed by Surface Plasmon Resonance (SPR) using an Biacore (Cytiva, T200). The Protein A chip was placed into the chip compartment, and the priming procedure was run to flush the system fluidics. The ligands, ActRIIA-hFc and ActRIIB-hFc, were each diluted to 2 μg/mL with running buffer HBS-ET⁺ and then injected sequentially onto the active flow cells (Fc2 and Fc4) of the Protein A chip at a flow rate of 10 μL/min to achieve capture levels of 75 RU and 125 RU, respectively. No ligand was captured on the reference flow cells (Fc1 and Fc3). The analytes were serially diluted in running buffer and then injected sequentially over flow cells Fc1–Fc4 at a flow rate of 30 μL/min, with an association time of 120 s and a dissociation time of 420 s. All association and dissociation steps were performed in running buffer. After each analyte concentration cycle, the chip was regenerated with Glycine 1.5 (10 mM glycine, pH 1.5) at a flow rate of 30 μL/min for 30 s to remove the captured ligand and any remaining analyte. Data analysis was performed using the Biacore T200 Evaluation Software, and the sensorgrams were fitted with a 1:1 binding kinetic model.

### 3.3 ELISA detects antibody binding to ActRIIA/ ActRIIB

The binding of in-house purified antibody to ActRIIA&B was assessed by ELISA. Recombinant ActRIIA (GM-87256RP, Genomeditech) or ActRIIB antigen (GM-84198RP, Genomeditech) was coated onto 96-well plates at 0.5 μg/mL overnight at 4°C. After blocking with BSA, serially diluted antibody was added and incubated. Bound antibody was detected using HRP-conjugated Mouse Anti-Human IgG Fc secondary antibody (Cat. No. A01854, GenScript) followed by TMB substrate; absorbance was read at 450 nm. EC₅₀ values were calculated by four-parameter logistic curve fitting using GraphPad Prism.

### 3.4 Flow cytometry assay of ActRII antibody binding activity

The binding of the antibody to ActRIIA or ActRIIB was evaluated by flow cytometry using *ACVR2A* or *ACVR2B* -overexpressing HEK 293 cells. Cells were harvested and washed with PBS containing 1% BSA-PBS. Approximately 1×10^6^ cells per sample were incubated with serial dilutions of the antibody in staining buffer for 60 minutes at 4°C. After washing twice with staining buffer, cells were incubated with FITC-conjugated Goat Anti-Human IgG Fc secondary antibody (Cat. No. ab97003, Abcam) for 60 minutes at 4°C in the dark. Cells were washed again and resuspended in PBS for analysis. Fluorescence intensity was measured using a FongCyte flow cytometer (ChallenBio, China). Data was analyzed using the device’s analysis software, and the media fluorescence intensity (MFI) was plotted against antibody concentration. EC_50_ values were calculated by four-parameter logistic curve fitting using GraphPad Prism.

### 3.5 Validation of ActRII mAb blockade of the SMAD signaling pathway

To validate the blocking effect of the ActRII antibody on the SMAD signaling pathway, we purchased HEK-Blue™ TGF-β Cells (InvivoGen, hkb-tgfbv2) and overexpressed ActRIIA and ActRIIB target proteins in these cells. During the experiment, normally cultured cells were seeded at 6 × 10⁵ cells/mL, 100 μL per well, and cultured overnight. The next day, the antibody was serially diluted 4-fold starting from 25 nM, incubated with the target cells for 1 hour, followed by addition of Activin A (Acro, ACA-H421b). The cells were then incubated in a CO₂ incubator for 6 hours. The supernatant was collected and assayed using Quanti-Blue solution (InvivoGen, rep-qbs), and the absorbance was read at 620 nm using a microplate reader. IC₅₀ values were calculated by four-parameter logistic curve fitting using GraphPad Prism.

### 3.6 Validation of the inhibitory effects of an ActRII mAb on ActRIIA and ActRIIB individual signaling pathways

The H_ActRIIA Reporter Cell Line (GM-C28074, Genomeditech) and the H_ActRIIB Reporter Cell Line (GM-C28076, Genomeditech) were purchased. These two cell lines overexpress ActRIIA or ActRIIB, respectively, while the corresponding ActRIIB or ActRIIA is knocked out. A luciferase reporter system was used for signaling pathway detection. In the experiment, cells were seeded at 1.5 × 10⁵ cells/mL, 100 μL per well, and cultured overnight. On the following day, the antibody was incubated with the target cells for 1 hour, followed by the addition of Activin A (ACRO, ACA-H421b) and further incubation for 6 hours at 37°C in a CO₂ incubator. After incubation, luciferase activity was detected by adding an equal volume of luciferase assay reagent (GM-040505C, GMBright One-Step, Genomeditech) to each well. Luminescence was measured using a multimode microplate reader (Infinite 200 PRO, Tecan). Data were analyzed using GraphPad Prism software. Luminescence intensity (relative light units, RLU) was plotted against antibody concentration, and the half-maximal inhibitory concentration (IC₅₀) was calculated by four-parameter logistic curve fitting.

### 3.7 293T cell non-specific binding assay

Target 293T cells were seeded into a well plate at a concentration of 1 × 10⁶ cells/mL with 100 μL per well. Monoclonal antibodies were incubated with the cells at 100 μg/mL (100 μL per well) for 1 h at 4 °C. After incubation, cells were washed with 200 μL of FACS buffer and centrifuged at 350 g for 5 min, and the supernatant was discarded. Subsequently, a Goat pAb to Human IgG (Delight 488) fluorescent secondary antibody (Abcam, AB97003) diluted 1:200 was added at 100 μL per well, and the cells were incubated for 1 h at 4 °C in the dark. Following incubation, cells were washed again with 200 μL of FACS buffer and centrifuged at 350 g for 5 min, and the supernatant was removed. Finally, cells were resuspended in 150 μL of FACS buffer per well, and the fluorescence signal intensity was detected and analyzed by flow cytometry to evaluate the non-specific binding level of the antibodies.

### 3.8 Tm and Tagg Analysis

Thermal stability was assessed using an Uncle system (Unchained Labs). Protein samples were heated from 25 °C to 90 °C at a ramp rate of 1.0 °C/min, with a 30 s incubation at 25 °C and a 30 s plate hold. The unfolding transition (Tm) was monitored by intrinsic fluorescence (350/330 nm ratio), and the aggregation onset temperature (Tagg) was determined by static light scattering (266 nm and 473 nm).

### 3.9 SEC Chromatography Conditions

The size exclusion chromatography (SEC) was performed on (aTSKgel G3000SWXL) column using a mobile phase consisting of 0.1 M Na₂SO₄ and 0.1 M phosphate buffer (pH 6.5) at a flow rate of 0.8 mL/min, with UV detection at 280 nm.

### 3.10 Animal study

#### Test article

Semaglutide used in the study was Ozempic (Novo Nordisk, 202502CJF2), and the monoclonal antibody was manufactured by Gene Universal (Anhui) Co., Ltd.

#### Diet and grouping

Male C57BL/6JGpt mice aged 5–6 weeks old (Strain NO.N000013) were purchased from GemPharmatech (Nanjing, China) and the study was performed according to the regulations and SOP of GemPharmatech LLC. Animals were group housed and maintained on a 12 h light/12 h dark cycle at standard temperature and humidity conditions and fed a high-fat diet *ad libitum* for 17-19 weeks prior to the start of study.

#### Pharmacodynamic assay

Mice were randomized into each groups (n=8 per/group) based on body weight and body fat percentage and were subcutaneously treated one of the following treatments: Antibody treatments were treated on days 0, 7, 14, and 21, animals received either 20 mg/kg each of antibody or Negative control (NC, commercially sourced PBS solution), semaglutide was dosed daily at 0.123 mg/kg, the combination group received the same regimen as the monotherapy groups. Body composition was assessed using a Low-field NMR body composition analyzer (Niumag Analytical, Suzhou), measured on days 0, 13, 20, and 27 of the study. Cumulative food intake was determined by manually weighing food from single cages twice a week. Body weight was measured twice a week. Mice were euthanized on days 28 of the study.

### 3.11 Statistical analysis

One- or two-way ANOVA followed by Sidak’s test for multiple comparisons (with repeated measures for time series data) was used in all studies. For comparison among multiple groups at a single time point, one-way ANOVA was performed. All tests used the software GraphPad Prism. Significance was defined as \**P* < 0.05, \*\**P* < 0.01, and \*\*\**P* < 0.001.

## 4 Discussion

The therapeutic landscape of obesity has been substantially transformed by incretin-based therapies, particularly glucagon-like peptide-1 receptor agonists (GLP-1RAs). Despite their remarkable efficacy in reducing body weight, accumulating clinical evidence indicates that a considerable proportion of weight loss achieved with GLP-1RA treatment consists of lean mass reduction. In clinical studies of semaglutide and tirzepatide, lean tissue has been reported to account for approximately 25–40% of total weight loss^[33]^. Because skeletal muscle is a major determinant of glucose disposal, energy expenditure, and physical function, excessive lean mass loss may compromise long-term metabolic health and potentially contribute to weight regain after treatment discontinuation. These observations have prompted increasing interest in therapeutic strategies that improve body composition rather than simply reducing body weight. An ideal anti-obesity intervention would maximize fat loss while preserving, or potentially increasing lean mass. The activin type II receptor pathway represents an attractive target for achieving this goal. ActRIIA and ActRIIB mediate signaling by myostatin, activins, and related TGF-β family ligands that negatively regulate skeletal muscle growth. Inhibition of this pathway has been shown to increase muscle mass and improve metabolic outcomes in both preclinical and clinical settings.

The clinical activity of bimagrumab established proof-of-concept for ActRII blockade as a body composition–modifying strategy. However, opportunities remain to improve potency, developability, and compatibility with current obesity therapies. In this context, we developed AB130-165, a novel dual ActRIIA/B-blocking antibody generated using AI-driven antibody engineering platforms, and evaluated its potential as a muscle-preserving adjunct to GLP-1–based weight management.

The development of AB130-165 illustrates the potential of AI-enabled antibody engineering for the discovery of functionally optimized biologics. Through iterative design and screening using the AlfaDAX platforms, we obtained variants with progressively more favorable predicted binding free energies (ΔG), which were positively correlated with experimentally measured affinities. The lead candidate, AB130-165, achieved sub-picomolar affinity for both ActRIIA and ActRIIB — representing more than a 1,000-fold and 10-fold improvement over the bimagrumab analog, respectively. Another variant, AB130-188, exhibited even more favorable predicted ΔG and correspondingly higher SPR-confirmed affinity, yet its cellular functional potency (IC₅₀) in the SMAD-responsive reporter system was inferior to that of AB130-165. Notably, while the physical binding affinity for ActRIIA (K_D_) surged by over three orders of magnitude, the cell-based functional blocking activity (IC₅₀) in the SMAD responsive reporter system experienced a more modest (∼10-fold) augmentation. This observation is consistent with previous findings^[34]^ in receptor pharmacology, where improvements in equilibrium binding affinity do not always translate proportionally into functional activity. Once receptor occupancy approaches saturation, additional gains in affinity may yield diminishing returns due to factors such as receptor turnover, ligand abundance, receptor clustering, and cellular signaling dynamics. The approximately ten-fold improvement in signaling blockade observed for AB130-165 therefore suggests that factors beyond affinity alone contribute to functional efficacy. Collectively, these findings highlight the importance of integrating structural and functional optimization during antibody development. The AlfaDAX platform successfully exploited this boundary condition: it did not merely chase affinity metrics, but optimized the functional interface spatial geometry to ensure maximum signal disruption *in situ* while using the ultra-low K_D_ to enforce prolonged receptor occupancy. Most current AI antibody design platforms — including RFdiffusion, Chai-2, JAM-2, and BoltzGen — remain primarily focused on generating high-affinity binders, with limited integration of functional or developability optimization. While these platforms have demonstrated impressive capabilities in *de novo* design and affinity maturation, their evaluation metrics are typically centered on binding detection rather than cellular function or biophysical liability filtering.

A recurring bottleneck for high-affinity biologics is the severe trade-off between functional potency and physical developability. Antibodies engineered for hyper-affinity often present high surface hydrophobicity, leading to self-aggregation, poor solubility, excessive viscosity at clinical concentrations—liabilities that complicate subcutaneous (SC) formulation development. AB130-165 successfully overcomes these biophysical limitations. *In silico* benchmarking via AlfaDAX predicted a favorable biophysical profile for AB130-165, maintaining low risk scores for aggregation propensity, viscosity, and immunogenicity, all comfortably below the critical commercial liability threshold of 1.0. These computational predictions were confirmed by experimental CMC analytics. AB130-165 showed good hydrophilicity, characterized by a Hydrophobic Interaction Chromatography (HIC) retention time of 20.7 minutes, matching the bimagrumab analog. Furthermore, SEC-HPLC analysis showed a monomeric purity of 97.17%, with a thermal stability profile (Tm = 69.30℃, Tagg = 66.45℃) that indicates a low risk of structural denaturation under physiological conditions, and supports stable long-term liquid formulation. FACS assays on HEK-293T cells demonstrated lower non-specific binding. . Combined with an acceptable transient expression yield in mammalian CHO cells (312.0 mg/L), these findings confirm that the computational mutation approach optimized the functional interface geometry while maintaining structural stability. The excellent biophysical and functional characteristics of AB130-165 make it amenable to a stable, high-concentration liquid formulation for low-volume, long-acting SC injection—a significant patient-compliance advantage over the cumbersome intravenous administration typically required in clinical settings.

The preclinical validation of AB130-165 in the C57BL/6J diet-induced obesity (DIO) mouse model provides clear proof of its best-in-class potential. Over a 27-day evaluation period, the combination of AB130-165 (20 mg/kg, QW) with semaglutide (0.123 mg/kg, QD) achieved a total body weight reduction of -33.4%, significantly outperforming the -25.5% drop in the bimagrumab combination cohort and the -24.3% in the semaglutide monotherapy group. An analysis of NMR data establishes that the deeper weight loss in the AB130-165 combination cohort was driven entirely by an accelerated mobilization of adipose tissue alongside the parallel preservation of lean body mass. AB130-165 combined with semaglutide reduced fat mass by 77.8%, compared with 65.0% for the bimagrumab analog combination and 46.4% for semaglutide alone. Semaglutide monotherapy resulted in a marked depletion of lean mass (−15.1%), confirming the muscle-wasting liabilities inherent to sustained incretin-driven calorie restriction, and co-administration of AB130-165 limited this lean tissue attrition to -9.3%. This translated into the lowest body fat percentage and the most favorable body composition profile among all treatment groups. The lean mass-to-body weight ratio reached 67.3% and fat mass-to-body weight ratio reached 12.6% in the AB130-165 combination group, compared with 62.3% lean, 17.4% fat in the bimagrumab analog combination group and 54.4% lean, 26.1% fat in semaglutide monotherapy group. The superior body composition remodeling achieved by the AB130-165–semaglutide combination may be attributable, at least in part, to the markedly higher affinity of AB130-165 for ActRIIA and its enhanced blockade of activin A signaling, resulting in more effective inhibition of the ActRII pathway during GLP-1RA-induced weight loss.

While the clinical-stage evaluation of related molecules like bimagrumab supports the metabolic validity of the ActRII pathway, the clinical advancement of a sub-picomolar dual antagonist like AB130-165 requires careful navigation of several biological and translational challenges:

Since ActRIIA and ActRIIB are widely expressed across peripheral tissues, particularly on the dense vascular endothelium and large skeletal muscle beds. AB130-165 is highly likely to undergo significant target-mediated drug disposition (TMDD). At lower clinical doses, the high peripheral receptor sink may rapidly clear the antibody from systemic circulation, resulting in non-linear elimination kinetics and a compressed initial half-life (*t*_1/2_). Characterizing the precise dose threshold required to saturate this peripheral receptor sink without causing systemic toxicity is a critical objective for upcoming non-human primate (NHP) pharmacokinetic studies. In addition, AB130-165 acts via a pan-blockade mechanism against both ActRIIA and ActRIIB receptor subtypes to maximize muscle preservation. However, as illustrated in the receptor activation pathways, these receptors are engaged by multiple ligands within the wider TGF-β superfamily, including GDF11, activin B, and various bone morphogenetic proteins (BMPs). GDF11 and activin signaling networks are involved in vital homeostatic processes beyond muscle tissue, including the regulation of late-stage erythropoiesis in the bone marrow and iron distribution cascades^[35]^. Prolonged, near-complete blockade of these pathways carries potential risks for hematological toxicities, such as alterations in red blood cell indices or unexpected vascular remodeling. Upcoming toxicology studies must prioritize close monitoring of these homeostatic parameters. Last, Rodent metabolic configurations and adipose-to-muscle mass distributions differ substantially from human physiology. The rapid, deep body composition remodeling observed in DIO mice over a 27-day testing protocol must be scaled cautiously when designing human clinical trials, which typically extend over 48 to 72 weeks. Future clinical investigations should combine non-invasive dual-energy X-ray absorptiometry (DXA) or magnetic resonance imaging (MRI) body tracking to establish whether the muscle-preserving effects seen in preclinical models translate into durable functional improvements, such as enhanced physical grip strength or increased aerobic capacity in human patients.

In conclusion, this study demonstrates the utility of the AI-driven AlfaDAX platforms for the rapid discovery and functional optimization of therapeutic antibodies targeting ActRIIA/B. Through AI-guided affinity maturation and developability optimization, AB130-165 achieved sub-picomolar target affinity alongside an approximately 10-fold increase in cellular functional blockade while maintaining favorable biophysical properties. In combination with semaglutide, AB130-165 produced superior body composition remodeling in DIO mice by enhancing fat loss while mitigating lean mass loss, supporting the concept of high-quality weight loss. These findings establish AB130-165 as a promising next-generation adjunct to GLP-1–based therapies and highlight the potential of AI-enabled antibody engineering to accelerate the development of biologics with improved therapeutic performance.

## Supplementary

**Supp Figure 1.**
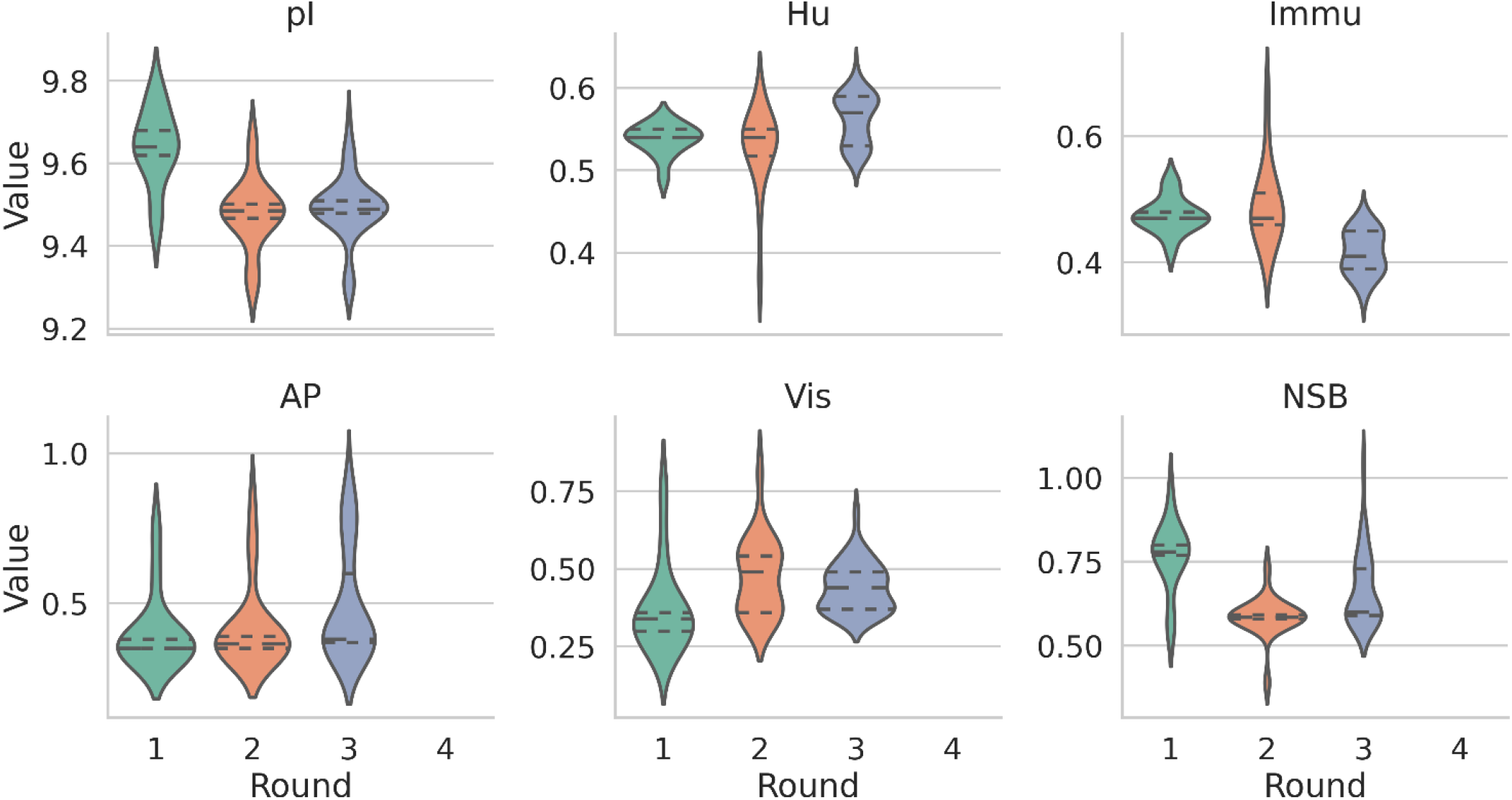
*In silico* developability metrics and confidence metrics of designed antibodies across iterative design rounds. Violin plots showing the distribution of key developability parameters for antibodies generated in rounds 1 to 4, as predicted by the AlfaDAX platform. Parameters include: (A) Isoelectric point (pI); (B) Humanization score (Hu); (C) Immunogenicity risk (Immu); (D) Aggregation propensity (AP); (E) Viscosity risk (Vis); (F) Non-specific binding (NSB). The central horizontal line represents the median, and the dashed lines indicate the quartiles. For Immu, AP, Vis, and NSB, scores ≤ 1 are considered low risk. Across all rounds, round 4 antibodies maintained favorable developability profiles, and all risk scores consistently below the threshold of 1.0, indicating low liabilities for immunogenicity, aggregation, viscosity, and non-specific binding.

